# A common haplotype in the EXO5 gene can impact its protein structure and dynamics and modulate genome stability and cancer progression

**DOI:** 10.1101/2024.11.22.623165

**Authors:** Fabio Mazza, Davide Dalfovo, Alessio Bartocci, Gianluca Lattanzi, Alessandro Romanel

## Abstract

Understanding the impact of common germline variants on protein structure, function, and disease progression is crucial in cancer research. This study presents a comprehensive analysis of the EXO5 gene, which encodes a DNA exonuclease involved in DNA repair previously associated with cancer susceptibility. We employed an integrated approach combining genomic and clinical data analysis, deep learning variant effect prediction, and molecular dynamics simulations to investigate the effects of common EXO5 haplotypes on protein structure, dynamics, and cancer outcomes. We characterized the haplotype structure of EXO5 across diverse human populations, identifying five common haplotypes, and studied their impact on EXO5 protein. Our analyses revealed significant structural and dynamic differences among the EXO5 haplotypes, particularly in their catalytic region. The L151P EXO5 protein variant exhibited the most substantial conformational changes, potentially disruptive for EXO5’s function and nuclear localization. Analysis of TCGA data showed that patients carrying L151P EXO5 had significantly shorter progression-free survival in prostate and pancreatic cancers, and exhibited increased genomic instability. This study highlights the strength of our methodology in uncovering the effects of common genetic variants on protein function and their implications for disease outcomes.

## Introduction

DNA double-strand breaks (DSBs) represent one of the most severe forms of genetic damage, posing a significant threat to genomic stability and cellular health. If left unrepaired or improperly addressed, DSBs can lead to genomic instability, potentially disrupting oncogens and tumor suppressor genes, thereby increasing cancer susceptibility and impacting cancer evolution (1)(2)(3). Two primary DNA damage response (DDR) pathways tackle DSBs: homology-directed repair (HDR) and non-homologous end joining (NHEJ). The choice between these pathways is governed by various factors, including cell cycle checkpoints and the activation of specific DNA repair genes. Germline genetic variants are the primary form of DNA polymorphism and variants in genes encoding proteins involved in DDR have been shown to contribute not only to cancer susceptibility but also to play a crucial role in determining treatment response and clinical outcomes (4)(5)(6). Notably, the process of DNA end resection, carried out by specialized exonucleases, plays a crucial role in determining the repair pathway (7)(8)(9).

Among those, EXO5 is a single-stranded DNA (ssDNA) exonuclease implicated in DNA repair, expressed by the *EXO5* gene. It is involved in homologous recombination following interstrand cross-links damage, specifically in the process of stalled DNA replication fork restart, where it performs DNA end resection (10)(11)(12). EXO5 loads at ssDNA ends, with 5’ to 3’ polarity enforced by Replication Protein A, and slides along ssDNA prior to the resection (10)(11 It plays a role in managing stalled replication forks, but its precise function in this process is not fully understood.

Ali et al (11) identified *EXO5* as a risk gene for prostate cancer (PCa). They reported multiple *EXO5* related germline Single Nucleotide Variants (SNPs) associated with PCa risk and demonstrated that knockout of *EXO5* gene leads to reduced HDR efficiency. Interestingly, the authors observed through in-vitro experiments that a mutation of residue 151 from Leucine to Proline, as a result of the common germline SNP rs35672330, causes a loss of EXO5 nuclease activity and of nuclear localization.

The role of EXO5 in cancer was further characterized in (12), where the authors observed that elevated EXO5 expression in tumors correlates with increased mutation loads and poor patient survival, suggesting that *EXO5* upregulation has oncogenic potential. The authors hypothesised that high EXO5 levels may contribute to mutations by potentially shunting repair of replication errors away from error-free homologous recombination and into more error-prone pathways. Structural data generated in the same study, indicates that the EXO5 channel where ssDNA inserts might contain a conditionally folded region, composed of an alpha-helix (*α*4) that unfolds upon DNA insertion. Further, the authors observed that without the presence of an iron-sulfur cluster bonded near the N-terminal of the protein, the nuclease function is mostly absent. The catalytic mechanism uses either a one-metal ion or a two-metal ion, and without availability of divalent cations such as magnesium ions the catalytic activity is also absent.

Although the role of EXO5 in cancer appears to be important for both tumor initiation and progression, comprehensive structural studies remain lacking. Notably, there has been no investigation into how common germline genetic variants might alter the structural conformation and dynamic properties of the EXO5 protein, which could have implications for its function in DNA repair pathways and its potential as a therapeutic target.

To address these knowledge gaps, here we utilized an integrated approach that combines the interrogation of large genomic datasets with deep learning predictions and Molecular Dynamics (MD) simulations. Initially, we assess and characterize common human EXO5 haplotypes using the ESM-1v pretrained protein language model (PLM) (13)(14). Following this, we conduct MD simulations to gain a mechanistic understanding of how the predicted fitness differences influence the protein’s structure and dynamics. Finally, we demonstrate the prognostic significance of the most impactful haplotype by analyzing cancer data from The Cancer Genome Atlas (TCGA).

Overall, our strategy enabled us to both explore how common human haplotypes affect the structural and dynamic properties of the EXO5 protein and identify potential genetic biomarkers for stratifying cancer patients according to distinct clinical outcomes.

## Materials and Methods

### Identification of transcript-specific haplotypes

Phased 1000 Genomes Project SNP genotypes were downloaded (www.internationalgenome.org/) and annotated using SnpEff v5.1d (15). Variants annotated as *Missense* were retrieved and combined with the human reference genome (hg38) to build the landscape of alleles for all protein-coding transcripts of EXO5 and a set of DDR genes of interest (16). Haplotype frequencies were then computed for each transcript across all 1000 Genomes Project individuals, and all haplotypes showing a frequency greater than 0.01 were selected.

### Calculation of haplotypes’ functional scores

To quantify the effect of germline SNPs within haplotypes on protein structures, we utilized the Evolutionary Scale Modeling ESM-1v protein language model (13). We use the Pseudo Log-Likelihood Ratio (PLLR) score (17) to quantify this impact, building on the work presented in (18):

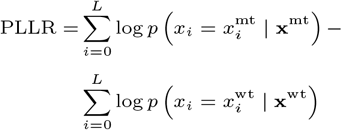

The PLLR score quantifies the difference between the model’s pseudo log-likelihood of a mutated protein sequence —specifically, one that incorporates changes introduced by haplotype-specific germline SNPs— and its corresponding wild-type protein sequence, by calculating their logarithmic difference. The log-likelihood, in this context, represents how likely a sequence is according to the model’s interpretation of protein *grammar*, which we estimate by summing the log-probabilities of amino acids as returned by the model’s output.

A positive PLLR score indicates that the variants make the sequence more plausible or less disruptive according to the model, while a negative PLLR score suggests a potentially harmful impact, implying that the variants may result in a sequence that is less likely to be functional or stable. More specifically, we compute the PLLR between SNP-containing sequences and their respective reference transcript sequence, using the ensemble of ESM-1v models 1 to 3. These correspond to the same ESM architecture with the same training data, but with different starting seeds for their training. We then consider the averages of the three resulting PLLR values for each non-reference sequence.

### Molecular Dynamics simulations

The impact of EXO5 haplotypes on the structural conformation and dynamics of the EXO5 protein was assessed through multi-replica, all-atom Molecular Dynamics (MD) simulations (19).

### Reconstruction and Modeling of EXO5 Protein Structures

The 7LW9 crystallographic structure of EXO5 (12) was obtained from the RCSB Protein Data Bank. The 7LW9 PDB file includes 255 modeled amino acids of the G172V variant of EXO5, with 118 residues missing, along with a 7-nucleotide single-stranded DNA molecule (TGAAGGG) and an iron-sulfur cluster [4Fe–4S]. The N-terminal (residues 1-68) and C-terminal (residues 358-373) regions were unmodeled due to their disordered nature. These regions have been shown in (12) not to contribute to EXO5 activity. All other missing residues (107-132), which include the conditionally-folded *α*-helix enclosing the channel over the ssDNA, were modelled superimposing the 7LW9 structure with the complete EXO5 structure we obtained using ColabFold (20)(In ColabFold, the AlphaFold2 model was used on the whole G172V EXO5 sequence, using the default MSA construction, no PDB templates, no relaxation of the structure and 12 recycles through the folding trunk of the model. Extraneous ions present in the 7LW9 PDB were then removed and the single *Mg*^2+^ ion resolved in the 7LWA was then copied to the final PDB after alignment of the two protein structures.

The resulting G172V EXO5 structure was then mutated to obtain the other EXO5 variants using PyMOL’s mutagenesis tool. The protonation of all structures was performed separately using PROPKA3 (22), via the playmolecule webserver (23). A doubly-protonated histidine (His117) was manually modified to have a single protonation state. The obtained EXO5 starting structure is shown in Figure 1.

**Fig. 1.**
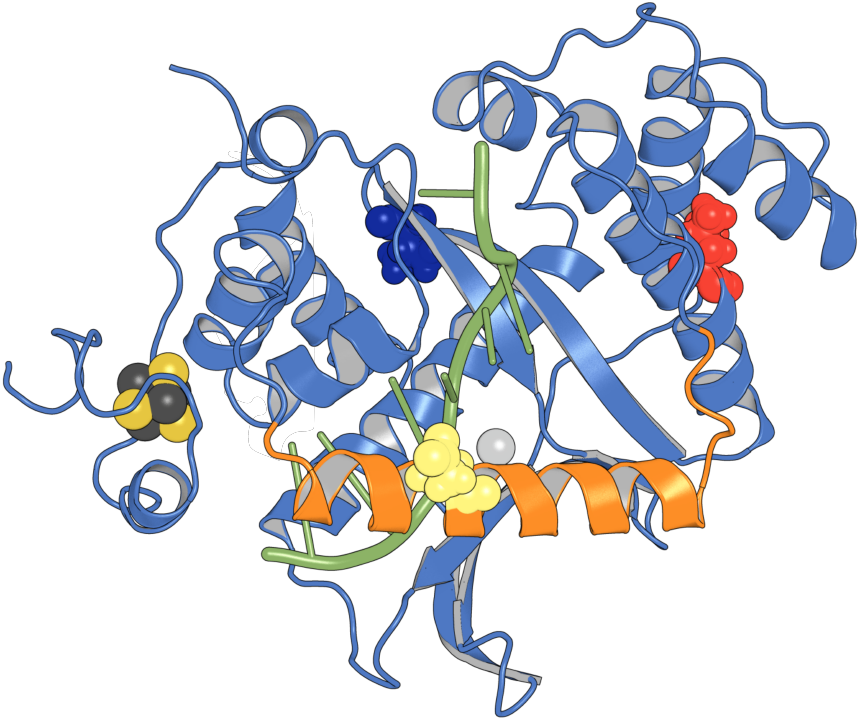
Illustration of the G172V EXO5 variant that was used as the starting structure of our simulations. In green the ssDNA, in light blue the structure taken from the 7LW9 PDB file, in orange the region extracted from the ColabFold prediction, which includes *α*4. The positions of the residues corresponding to the germline SNPs we have considered are marked with a sphere representation for the whole aminoacid: blue for G172, red for L151 and yellow for D115. The catalytic Mg^2+^ ion is represented as a grey sphere, and the [4Fe–4S] iron-sulphur cluster with yellow and dark grey spheres for sulphur and iron atoms, respectively.

The protein was described by the ff14SB Amber force field (24), in conjunction with the parmBSC1 force field for the ssDNA (25). The Li-Merz parameters were used for ionic species (potassium chloride and magnesium) (26). The iron-sulfur cluster was modeled based on the DFT-optimized geometry detailed in (27), specifically in its reduced state *A*_2+_. Four covalent bonds between the *Fe* atoms of the iron-sulphur cluster and protein cysteines Cys92, Cys356, Cys359 and Cys365 were finally added using AmberTools23 (28).

### Simulation Protocol

The EXO5 variants’ structures were used to create distinct simulation systems. Each system was solvated in water with a 15 Å buffer distance, and 187 potassium ions and 169 chloride ions were added to reach a 150 mM KCl concentration as well as ensuring the system’s overall charge neutrality.

MD simulations were carried out with Gromacs 2023.3 (29). All systems underwent energy minimization employing the steepest descent algorithm until a maximum force of 750 kJ mol^−1^ nm^−1^ was reached. This was followed by a heating phase of 50 ps, from 0 to 310K. The solvent was thus thermalized in the canonical (NVT) ensemble at T = 310 K for 250 ps, employing the V-rescale thermostat (30) (*τ*_*T*_ coupling constant of 0.2 ps) with an integration timestep of 1 fs. Then, an equilibration step of the solvent in the isothermal-isobaric ensemble (NPT) at P = 1 bar was carried out for 1 ns, *via* the C-rescale barostat (31) (*τ*_*P*_ = 2.0 ps, compressibility of 4.5 × 10^−5^ bar^−1^). During both stages, position restraints were applied to the magnesium ion, as well as to all heavy atoms of the protein and DNA, using force constants of 1000 kJ mol^−1^ nm^−2^. A final unrestrained NPT equilibration was performed for 2 ns with a time step of 2 fs.

For all systems, a 4 µs-long trajectory was produced, after which the first 960 ns were cut. The frames at 960 ns were used to start (with randomized velocities) other 2 independent replicas of each system. Ultimately, 3 production replicates of 3 µs each (in total 9 µs per system) were obtained for each EXO5 variant.

MD equilibration and production stages in the NPT ensemble were carried out using an integration time step *t*_*step*_ of 2 fs, the v-rescale thermostat with *τ*_*T*_ = 0.2 ps, T = 310 K, and the C-rescale barostat with *τ*_*P*_ = 2 ps at 1 bar. For the solvent, including all ions present in the system, and the solute, separated temperature couplings were used. Electrostatic interactions were treated using the PME method (32)(33) a PME order of 4. A cutoff of 1.1 nm was employed for both electrostatic and Van der Waals interactions. Each bond involving hydrogen was constrained with the LINCS algorithm (34)(35). Dispersion corrections for energy and pressure were applied using the dispcorr=EnerPres GROMACS parameter, as recommended for simulations employing Amber force fields to account for long-range Van der Waals interactions.

### MD trajectories analysis

The evolution of the EXO5 protein variants’ secondary structure elements, particularly in the region surrounding the *α*4 helix, was assessed using the MDAnalysis implementation of the PyDSSP algorithm (36)(37). This is a simplified implementation of the original DSSP algorithm (38), where *β*-bulges are determined as loops instead of *β*-strands. Hydrogen positions were explicitly parsed from the trajectories instead of guessed.

MDAnalysis was also used to calculate the Root Mean Square Deviation (RMSD) and Root Mean Square Fluctuations (RMSF) over the trajectories, with a sampling frequency of 200 ps for both RMSD and RMSF. The structures were aligned to the protein core, which excludes *α*4 because of its high mobility. The reference structure for RMSD was the PDB structure, while for the RMSF the average frame of the equilibrated trajectories was used. Different replicas were treated separately, averaging over the final results in the case of RMSF.

When an uncertainty on average values is reported, it represents the standard error of the mean, calculated based on the averages across replicas. The variance of the average estimate for each replica was derived using the Moving Block Bootstrap method applied to all frames within that replica, unless otherwise specified (39). The block size was determined by estimating the autocorrelation time of the relevant time series as the time-lag required for the autocorrelation function to reach the value *e*^−1^.

### Protein Structure Network and Residue Correlation Network

In order to characterize the protein structure networks of our systems we process the MD trajectories using PyInteraph 2 (40)(47). In this framework, the protein is described as a graph, with residues as nodes and edges defined by noncovalent interactions, which in the case of PyInteraph 2 are characterized with user-defined geometric criteria corresponding to different chemical interactions. We defined the edges using atomic contacts between side chains, using the PyInteraph default residue and atomic interaction definitions for hydrogen bonds, salt bridges and hydrophobic interactions. We further added to the residue selection for hydrogen bonds the DNA and the *Mg*^2+^ ion.

The network was then generated from the equilibrated trajectories, excluding the first 960 ns of the first replica, with a frequency of 100 ps. The resulting adjacency matrices were weighted based on the occurrence of the interactions, i.e. the fraction of frames where the contact is detected divided by the total number of frames. The networks for each trajectory were then combined by taking, for each edge, the maximum occurrence value among the three types of interaction. A set of unweighted networks was also extracted from the weighted ones, using an occurrence cutoff of 20%, in line with previous literature (40).

We then used the Jaccard similarity coefficient to evaluate the similarity across the different, unweighted networks. This coefficient was calculated by dividing, among all pairs of trajectories, the number of common edges by the the number of edges present in either one of the two networks.

### Dimensionality Reduction and Clustering

With the aim of comparing the essential dynamics of EXO5 in the simulated trajectories, we performed Principal Component Analysis (PCA) and computed the Root Mean Square Inner Product (RMSIP) of the resulting essential subspaces (42)(43)(44). Different replicas were analyzed separately, allowing an assessment of the convergence of the simulations along with the comparison between the different EXO5 haplotypes.

PCA was performed on the *α*-carbon positions of the protein trajectories. Prior to performing the PCA, the trajectories were aligned via the alpha-carbons of the first equilibrated frame of G172V EXO5, excluding residues 107-131. A frame every 0.5 ns was analysed, resulting in about 30000 configurations per trajectory. The first 30 principal components of each trajectory were then used to compute the RMSIP. Both the PCA and the RMSIP calculation were performed using MDAnalysis.

The protein conformations were also clustered using the Advanced Density Peaks Clustering algorithm (45) implemented in the DADApy package (46). Instead of using the 3D coordinates of the protein backbone, the *ϕ* and *ψ* backbone dihedrals were used as input, following previous research indicating that distance measures based on backbone dihedrals are more effective for clustering purposes (47). Protein dihedrals of the *α*4 region spanning residues 107-131 and the set of all other protein dihedrals were clustered separately, respectively with a minimum uncertainty for the cluster separation of *Z* = 5 and *Z* = 3. This separation was chosen based on preliminary analyses that showed most of the structural diversity between different trajectories was driven by the residues 107-131. Clustering was performed on the concatenated EXO5 trajectories, with a total of 38000 frames.

### Survival analysis and genomic instability analysis using TCGA data

Genotype and ancestry data of 8,535 TCGA cancer patients were retrieved from (48) where unphased genotypes for exonic SNPs were computed using (49) and ancestry information was retrieved by means of (50). Genotypes were then phased with SHAPEIT v2 (51) to infer haplotype structure using 1000 Genomes Project genotype data as reference panel. EXO5 haplotypes were then used to perform survival analysis using TCGA cancer survival data (52). Overall Survival (OS) and Progression-Free Interval (PFI) were considered in our analysis. Patients were stratified based on presence of specific haplotypes. Kaplan-Meier survival curves were generated, and Cox proportional hazards regression models were applied, adjusting for ancestry information. For prostate cancer, a significance threshold of p *<* 0.05 was used. For the extended analysis across other cancer types, only those with more than 100 patients were included, and multiple hypothesis correction was applied using a q-value threshold of 0.05. Survival analyses were performed using the survival R package (53).

Genomic instability analysis was conducted using data from (54). Specifically, weighted genome instability index values were obtained and analyzed for 310 patients with prostate or pancreatic cancer. Patients were stratified based on the presence of at least one allele carrying a specific EXO5 haplotype. Individuals with significant aneuploidy—defined by an allele-specific aneuploidy score below 1.5 or above 2.5—were excluded.

## Results

### Haplotype structure of EXO5 gene across human populations

To characterize the haplotype structure of the *EXO5* gene across diverse human populations, we utilized genome-wide phased genotype data from approximately 2,000 individuals obtained from the 1000 Genomes Project. We focused on identifying all missense SNPs and INDELs within the coding regions of all annotated *EXO5* gene transcripts, with the aim of focusing on common germline variants that could potentially alter the functional properties of the translated protein. Notably, although ENSEMBL v111 reports multiple transcripts for the *EXO5* gene, the coding sequence remains uniformly conserved within the final exon across all transcripts. Indeed, all transcripts are annotated as encoding the same protein, Q9H790, which consists of 373 amino acids.

As shown in Figure 2, we identified five common haplotypes of the *EXO5* gene, each with a frequency exceeding 1% across all individuals analyzed (around 4,000 haplotypes), collectively accounting for over 98% of the population. *Haplotype 1*, here also referred to as the wild-type (WT), corresponds to the reference human genome sequence (hg38) and was observed with a frequency of 40.4%. *Haplotype 2*, with a frequency of 42.3%, resulted as the most prevalent haplotype and was characterized by the presence of the alternative allele at SNP rs11208299. *Haplotype 3*, which carries the alternative alleles at both rs11208299 and rs1134586 SNPs, follows with a frequency of 10.6%. *Haplotype 4*, associated with the alternative allele at SNP rs35672330, was observed with a frequency of 2%, while *haplotype 5*, which includes the small deletion rs35672330, occurred with a frequency of 2.8%.

**Fig. 2.**
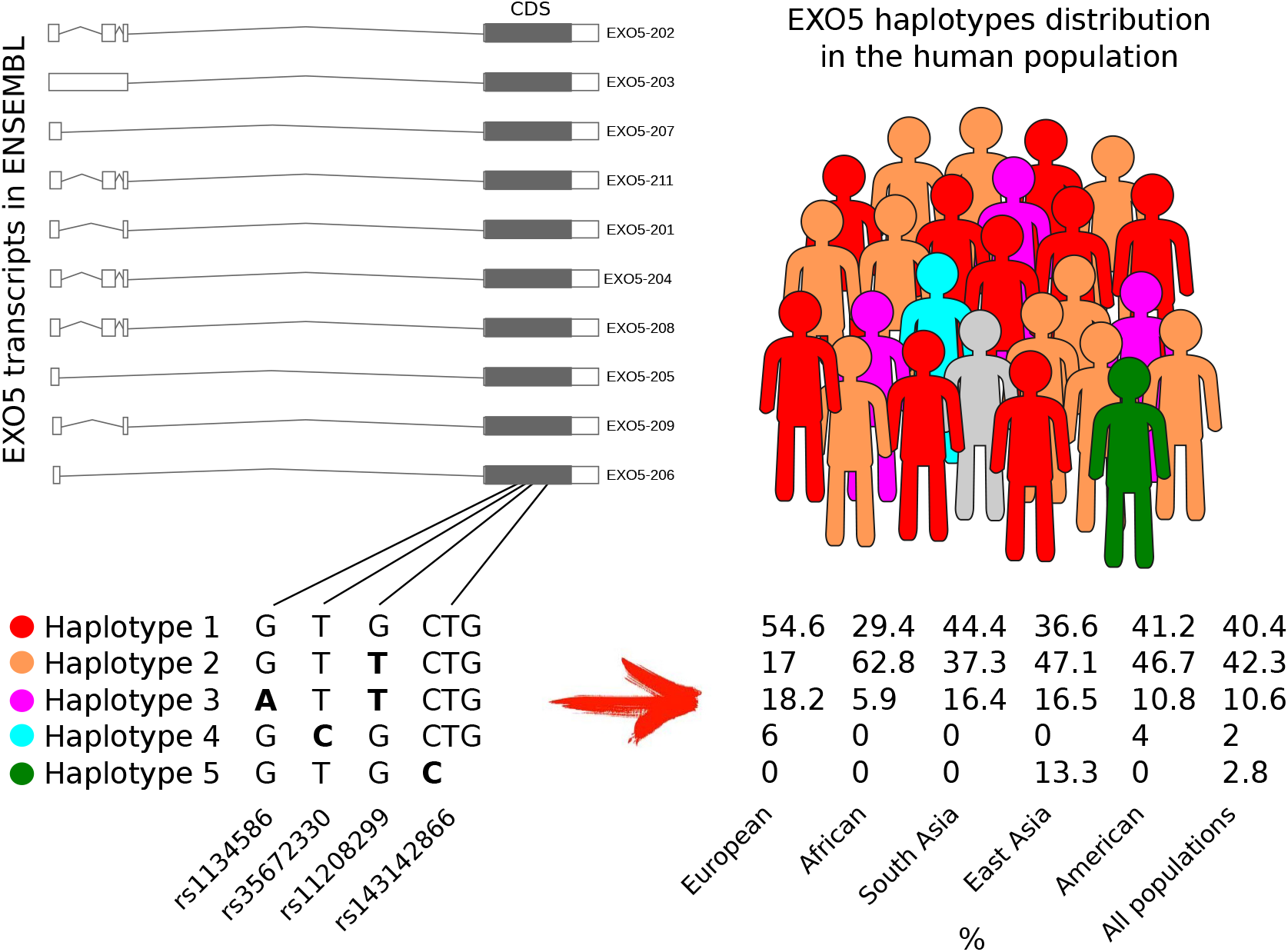
EXO5 haplotype structure across major human populations. On the left, the annotated EXO5 transcripts are depicted, along with their shared coding sequence (CDS), highlighting the position of the coding SNPs that define the common EXO5 haplotypes. On the right, the distribution of these EXO5 haplotypes is shown across the global human population and across the major populations.

Interestingly, the distribution of EXO5 haplotypes varied significantly across different populations. The WT haplotype was predominantly enriched in the European population, whereas *haplotype 2* was more frequently observed in African individuals. Despite its similarity to *haplotype 2, haplotype 3* exhibited a distribution pattern closer to that of the WT. Notably, *haplotype 4* was found only in Europeans and admixed Americans, while *haplotype 5* was exclusively observed in the East Asian population.

Overall, the haplotype structure of the EXO5 gene was heterogeneous both across individuals and populations and predominantly consisting of SNPs’ patterns.

### Quantification of common human haplotypes impact on EXO5 protein structure and function

Although the germline variants defining the common EXO5 haplotypes we identified are annotated as benign in ClinVar (55), we reasoned that recent advances in deep learning approaches could help us screen for any potential, even subtle, functional impact these haplotypes might have on the EXO5 protein.

In particular, to quantify the impact of missense variant patterns on the structure and function of the EXO5 protein, we employed the ESM-1v protein language model (13), notable for capturing long-range patterns and features essential for protein function and stability, making it particularly effective in assessing the impact of multiple alterations in amino acid sequences (14).

We focused our analysis on *haplotypes 1* through *4*, as these exhibit distinct SNP patterns and are represented across multiple major populations. First, we generated all EXO5 amino acid sequences incorporating the modifications caused by these SNPs. We then applied the ESM-1v model to calculate the Posterior Log-Likelihood Ratio (PLLR), a functional score that quantifies the impact of haplotype SNPs at the protein level by comparing the model’s log-likelihood of the SNP-altered amino acid sequence to that of the WT sequence.

Compared to the WT amino acid sequence of *haplotype 1*, the sequences for *haplotypes 2* and *3*, which are characterized by the G172V and G172V+D115N amino acid changes, respectively, showed positive scores of 5.05 and 4.51. In contrast, the *haplotype 4* sequence, characterized by the L151P amino acid change, presented a negative score of -2.74. These PLLR scores suggest that the G172V and G172V+D115N variants of the EXO5 protein may be more *favorable* compared to the WT protein, while the L151P variant appears to be significantly *unfavorable*.

To better assess the significance of the score variability observed for EXO5, we extended our analysis to a set of 251 genes associated with DNA damage response and repair, identified from the literature (16). Specifically, we retrieved missense SNPs across these genes, determined the common haplotypes for all gene transcripts, and calculated the PLLR scores of these sequences. The resulting score distribution, which includes the 174 genes observed to have multiple common haplotypes, is shown in Figure 3A. As shown, although the majority of PLLR scores clustered around zero (indicating haplotypes with no significant impact compared to the corresponding WT protein sequence), over 15% of haplotypes exhibited scores with an absolute value greater than 1. Furthermore, among the 721 transcripts with multiple haplotypes observed across all populations, 90 (12.5%) displayed a score variability (calculated as maximum difference between transcript specific PLLR scores) greater than 1 (Figure 3B). Notably, the EXO5 score variability of 7.8 was situated at the upper tail of this distribution, second only to RAD51B, which showed a maximum variability of 8.5. Additionally, an analysis of cancer somatic mutational profiles from the TCGA dataset (via cBioPortal) revealed that among the genes with haplotype variability greater than 2, several genes - including *EXO5, RAD51B, NUDT15, SWSAP1, POLM*, and *CLK2* - are recurrently altered in at least 2% of patients across various cancer types, based on data from approximately 11,000 TCGA patients.

**Fig. 3.**
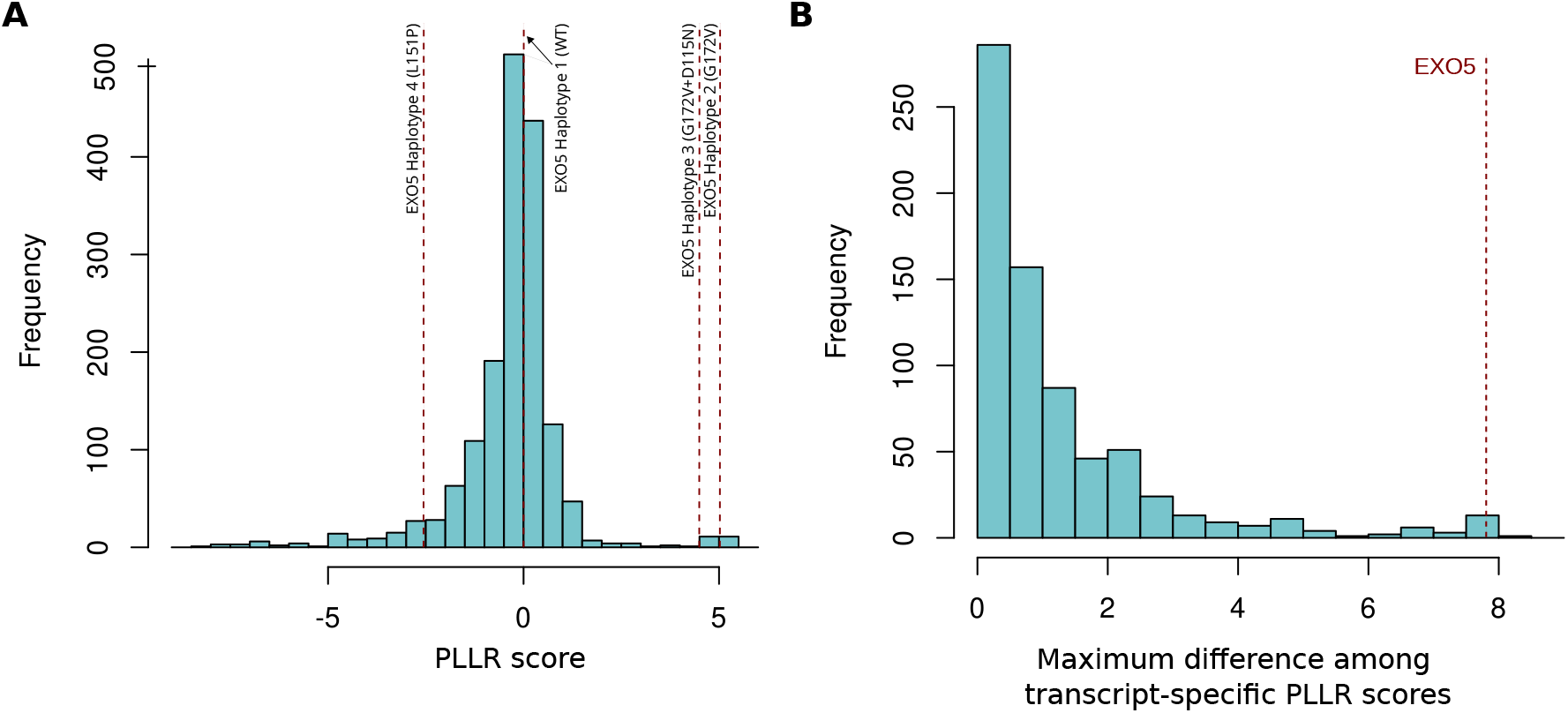
(A) Distribution of the PLLR scores calculated across all DNA damage response and repair genes. (B) Distribution of the maximum difference of scores between variants inside each of the 174 genes with at least one common SNP.

Overall, this data strongly supports the hypothesis that the various EXO5 haplotypes, though common in the population, result in different EXO5 protein variants that may have distinct structures and functional properties, and could potentially interact with cancer somatic alterations.

### Mechanistic effects of common EXO5 haplotypes on protein structure, stability, and dynamics

Considering the high variability of haplotype PLLR scores we observed for EXO5 protein variants and the availability of the resolved structure for G172V EXO5 in PDB (7LW9), we investigated mechanistically the effects of different EXO5 haplotypes using MD simulations. Starting from the available G172V EXO5 structure, we generated all EXO5 protein variants corresponding to SNP-based haplotypes. The initial structure, corresponding to *haplotype 2*, was thus reverted to the wild-type (*haplotype 1*) by mutating Val172 to Gly172. From the WT EXO5 structure, L151P EXO5 was then obtained (*haplotype 4*), while adding the D115N substitution to the G172V EXO5 structure resulted in the protein associated with (*haplotype 3*).

We then performed MD simulations of all systems in the presence of a ssDNA, in order to better characterize the native structure and dynamics of the active state of the nuclease. Such computational approach is well suited to catch the conformational dynamics of the DNA strand (56). Its binding region, being directly influenced by residues’ substitutions, is especially sensitive to haplotypes. Protein’s activity, in fact, can be allosterically modulated by the binding with biological molecules (57; 58; 59), and be affected by mutations (60).

None of the haplotypes produced a global destabilization of the protein tertiary structure, but significant differences, consistent among different replicas, were apparent in the structure and dynamics of the region spanning residues 106 to 133, comprising the *α*4 helix and the two disordered arms connecting it to the core of the protein (Figure 4A).

**Fig. 4.**
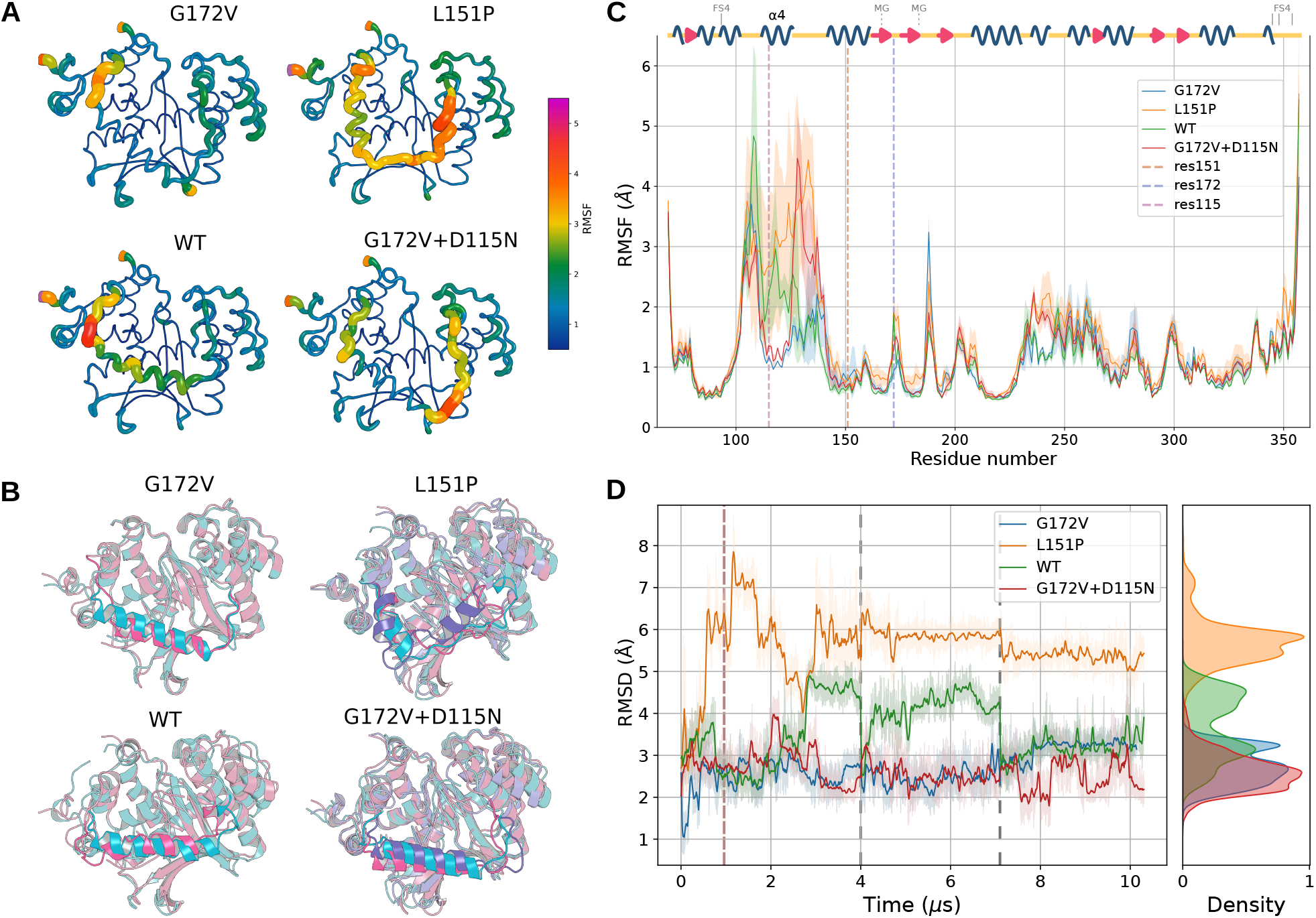
(A) Average conformation of EXO5 structures across the equilibrated trajectories. The size and color of the local representation are given by the RMSF for each residue. (B) Representative EXO5 structures with the ssDNA ligand. Each color represents one cluster center as found by clustering the backbone dihedrals of residues 106-133 obtained from the equilibrated trajectory. Only clusters containing at least 10% of the total frames are shown. Residues 106-133, which include the *α*4 helix, are highlighted in front of the protein structures. Full clustering results are reported in figure S4. (C) Average RMSF of EXO5 calculated separately for all replicas, with confidence intervals calculated as the standard error of the mean. On top of the RMSF plot, the secondary structure of G172V EXO5 is shown, as computed on the whole MD trajectory and then filtered to show as structured residues only the ones with the same assigned type for more than 70% of the frames. The sequence positions of residues coordinating the magnesium ion or forming the coordinate bonds with the iron-sulphur cluster are also pointed out as dashed lines. (D) RMSD of the *α*4 region (residues 107-131) across the whole trajectory. Lighter shades are the raw data, calculated every 0.5 ns, while the darker lines are a moving average with a 40 ns uniform window. The dashed red line indicates the chosen equilibration cutoff, while the dashed gray lines separate different replicas.

The *α*4 helix, resolved in (12) for EXO5 with no ligands, was absent in the DNA-bound G172V EXO5 structure, suggesting it may be a conditionally folded region or exhibit higher mobility in the presence of ssDNA. AlphaFold 2 (for G172V EXO5) predicts this region to form a 19-residue-long helix spanning residues 109 to 127. This helix, along with the disordered linkers, is proposed to form a channel for ssDNA. In contrast to the DNA-free structure of 7LW7, the position and length of the *α*4 helix in the predicted structure differ in order to accommodate the ssDNA.

### Alpha helix 4 undergoes big conformational changes in L151P and WT EXO5

Our simulations showed an overall stability of the *α*4 helix conformation for the G172V EXO5 variant, with an average 5.5 Å RMSD with respect to the starting EXO5 structure (Figure S1) and a 2.5 Å RMSD when aligning the 106-133 region alone (Figure 4D). This discrepancy is the result of a component of rigid motion of the *α*4 region. Indeed, in G172V EXO5 simulations, the disordered linker corresponding to residues 128-138 relaxed towards *α*5, pulling toward this region and thus towards the DNA channel the N-terminal part of *α*4 as well. Despite this displacement, which is common to all the simulated EXO5 variants, the local conformation predicted by Alphafold for G172V EXO5 is overall confirmed. This stability is also supported by RSMF values (Figure 4C) and by an analysis of the region’s secondary structure performed with PyDSSP (Figure S3), which shows an average difference of 20% between the secondary structure assignment during the (equilibrated) simulation and in the starting PDB structure, mostly due to the formation of transient secondary structures within the linkers adjacent to *α*4 (Figure S2B).

Notably, the addition of the D115N SNP does not disrupt the alpha-helix 4 conformation (Figure 4A), with an average RMSD around 2.5 Å and a secondary structure compatible with G172V EXO5 (Figure 4B and Figure S2), but results in higher RSMF values in residues 124-135 (Figure 4C), the linker adjacent to *α*5 and *α*7, suggesting possible changes in the allosteric interactions between *α*4 and other EXO5 regions. When focusing on L151P EXO5 protein variant, *α*4 and the surrounding linkers were observed undergoing substantial structural rearrangements, higher RMSF values, and changes in the secondary structure (Figure 4). This conformational change is clearly visible in Figures 4A and 4B, where the average and the most common conformations of L151P EXO5 show a substantial loss of the original *α*4 helix structure, replaced by a disordered region that spans the residues between Trp116 and Trp137. The only persisting stable alpha-helical structure is formed between residues Pro108 and Thr115, partially replacing the short disordered linker in the starting structure. This reorganization is reflected in a lower fraction of ordered residues in the *α*4 region of L151P EXO5 (ordered fraction 0.422 ± 0.033) when compared to WT EXO5 (ordered fraction 0.534±0.046) and to G172V EXO5 (0.524±0.043). The fraction of secondary structure assignment changes compared to the starting structure, shown in Figure S3, as well as the RMSD in

Figure 4D, highlight the strong deviation of the *α*4 region in L151P EXO5.

Finally, when analyzing the WT EXO5 variant, we observed a high variance between replicas in the RMSD and secondary structure metrics of the *α*4 region. For the first 2.5 µs of the first replica, the dynamics is similar to the G172V EXO5. At 2.5 µs, as well as shortly after the beginning of the second replica, the alpha-helix exhibits a loss of stability corresponding to higher RMSD values and changes in the secondary structure (Figure 4C and 4D). However, even during those time spans we did not observe such a drastic conformational change as in L151P EXO5. As indicated by the RMSF values (Figure 4C), and opposed to what happens in L151P EXO5, it is the short disordered linker composed by residues 103-109 that manifests higher RMSF values, with the long linker on the opposite side maintaining RMSF values below 2 Å, compatible with the ones of the G172V EXO5. The integrity of the *α*4 helix exhibits instability, falling in between the behaviors observed in EXO5 L151P and G172V variants.

Overall, L151P EXO5 exhibits the highest mobility and conformational variety in the *α*4 region, resulting in unique, highly disordered, conformations, as evidenced in Figure 4B. On the other hand, the two EXO5 variants with the G172V change present the greatest similarity to the predicted initial conformation and to each other. WT EXO5 shows intermediate behavior, adopting unique conformations with less variability and maintaining a secondary structure that more closely resembles the one predicted by AlphaFold, compared to L151P EXO5.

These results can also be observed when clustering the dihedral angles in the region (Figure S4). Notably, apart from a minor fraction of frames where G172V and G172V+D115N EXO5 variants share the same cluster, each remaining cluster is predominantly associated with a single EXO5 variant. In particular, the L151P EXO5 variant displays a greater diversity of clusters, with many clusters containing frames unique to only one of the three replicas.

Remarkably, the stability of *α*4, part of the catalytic region of EXO5, correlates with the predicted fitness scores produced by the ESM PLM analysis, even if none of the mutated residues that influence the scores are structurally close to *α*4, except for Asn115.

### Structural and dynamical consequences of the L151P substitution propagate from a local kink in helix α5

It is known that proline substitutions limit the flexibility of their peptide bonds, disrupt the local hydrogen bond network and introduce a kink when found in the middle of alpha helices. Indeed the most direct effect of the L151P substitution in EXO5 is the formation of a kink in *α*5, as revealed by the changes in the angle formed by its extremes, with Leu151 as its vertex (Figure 5A). Although the kink formation occurs in a time scale of microseconds, possibly indicating a suboptimal sampling with our simulation time, the angle sharply decreases in all L151P EXO5 replica, reaching values below 120° (average 133.0° ± 3.3°), compared to an average of 144.3° ± 1.8° in WT EXO5 (Figure 5B).

**Fig. 5.**
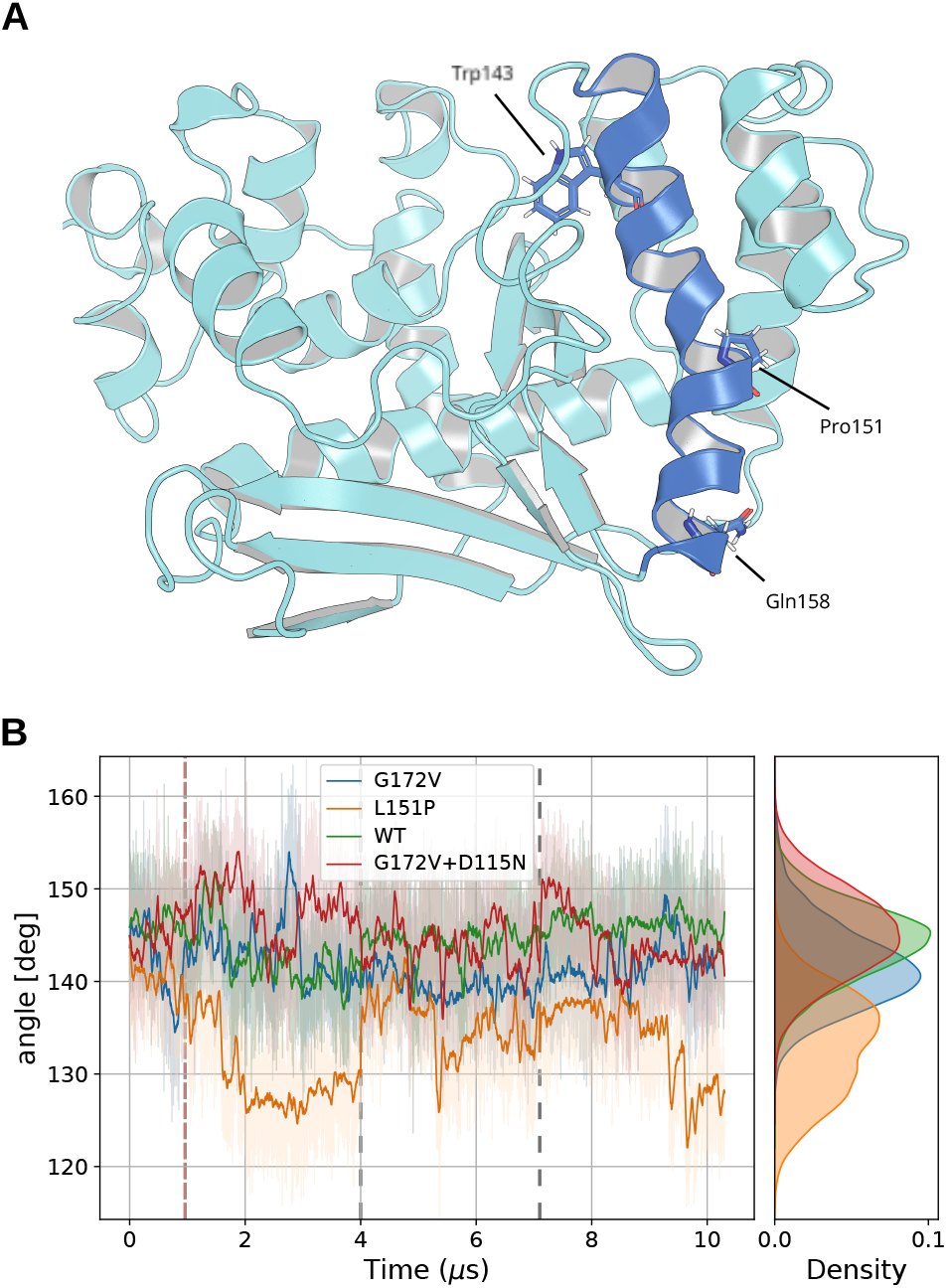
Illustration of a representative structure of L151P EXO5. *α*5 is highlighted in blue, and the Pro151 residue is highlighted, along with the residues used to compute the angle formed by the *α*-helix. The trajectory frame was chosen as the center of the biggest cluster along all replica of the L151P EXO5 MD trajectory (A). Angle formed by Trp143, Pro151 and Gln158 (B). Lighter shades are the raw data, calculated every 0.5 ns, while the darker lines are a moving average with a 40 ns uniform window. The dashed red line indicates the chosen equilibration cutoff, while the dashed grey lines separate different replicas.

This kink in *α*5 likely contributes to the destabilization of *α*4, a hypothesis further supported by the increased RMSF observed in the L151P EXO5 variant (Figure 3C), particularly within the linker region connecting these two alpha-helices. This strongly suggests that the stability of *α*4 is likely compromised by the L151P substitution through an allosteric pathway that propagates across, or through, the linker between these regions.

The conformational changes in the region comprising *α*4 and its surrounding disordered loops are also found to allosterically propagate the effects of the L151P substitution to the opposite side of the protein. This region is particularly interesting because it contains a sequence motif (KTRRRPMLPLEAQKKK, residues 198-213), which is classified with a high confidence by INSP (61) as a Nuclear Localization Signal (NLS). Especially relevant are the arginine and lysine triplets, which are known targets of the Importin class of transport proteins. The analysis of the network of interactions made with PyInteraph (Figure 6A) shows that in L151P EXO5, the middle arginine of the polybasic motif (R201) forms a stable salt bridge with D115, an aspartate that forms a transient alpha helix with the surrounding residues only in L151P EXO5 (Figure 6B). In 70.5% of the equilibrated frames (Figure 6A), the charged side chains of R201 and D115 are less than 4.5 Å distant. In Figure 6C it can be seen that this interaction occurs in all simulation replicas, while for WT EXO5 it is a transient interaction, with R201 and D115 isolated for most of the 9 µs trajectories. This interaction is completely absent in G172V and G172V+D115N EXO5 (see also Figure S5).

**Fig. 6.**
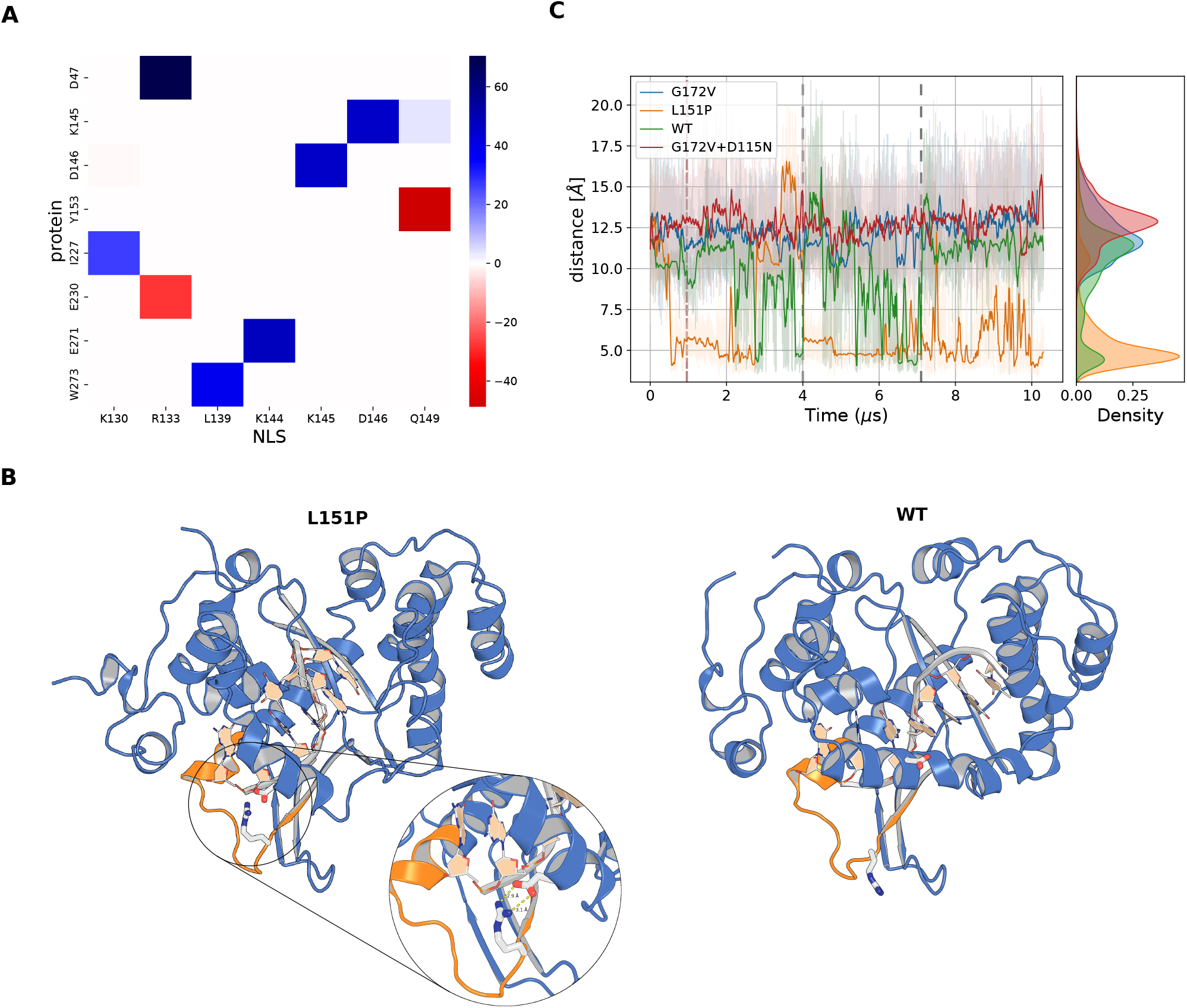
(A) Difference in the occurrence of contacts between the nuclear localization sequence and the rest of the protein, in the MD trajectory of L151P EXO5 vs the trajectory of G172V EXO5; (B) Illustration of the NLS (orange), with Arg201 and Asp115 represented with blue and white sticks and red and white sticks, respectively. The wild-type system is shown below for comparison. The structures shown are the representative frames of the respective system’s trajectories, as found with clustering, by identifying the frame with the highest logarithmic density inside the largest cluster. (C) Distance between Arg201 (zeta carbon) and Asp115 (gamma carbon) over time. Lighter shades are the raw data, calculated every 0.5 ns, while the darker lines are a moving average with a 40 ns uniform window. The dashed red line indicates the chosen equilibration cutoff, while the dashed grey lines separate different replicas.

Additionally, on the lysine triplet, K212 of L151P EXO5 forms a salt bridge with E339 (63.0% of occurrence) and K213 with D214, part of the same NLS motif. These contacts are only transient (14%-21% occurrence for K212-E339) in all other EXO5 simulations (see Figure S5). These newly formed contacts do not cause a large, observable, conformational change in the NLS backbone, but they seem to result in a reduced mobility of the disordered loop instead. This is indicated by a much longer tail in the autocorrelation function of the NLS RMSD (Figure S6B), which corresponds to an estimated autocorrelation time one order of magnitude higher compared to the autocorrelation time for all replicas of G172V, WT and G172V+D115N EXO5. Not only the RRR and KKK motifs recognition by an importin could be hindered by a high affinity to D47 and E271 in L151P EXO5, but if this recognition is dependent on a specific conformation of the loop, the higher autocorrelation time of the NLS loop indicates that a lower loop mobility could considerably slow down the process.

### Global structural and dynamic differences among EXO5 variants

The similarity analysis of the entire EXO5 structures reveals a pattern consistent with that observed in the *α*4 region.

The unweighted PSNs derived from the equilibrated trajectories of the G172V and G172V+D115N EXO5 variants show the highest similarity, as reflected in their Jaccard scores (Figure 7A). WT EXO5 also has comparable similarity to G172V EXO5, but slightly lower compared to G172V+D115N EXO5. L151P is instead an outlier, with an average Jaccard similarity of 0.5 compared to G172V EXO5 and 0.55 compared to WT EXO5. While the semi-disordered region in WT EXO5 is more similar to L151P EXO5 than to other variants, the overall interaction network of WT EXO5 aligns more closely with the G172V variants, indicating that its residue interaction network is less affected by the instability of *α*4. Both G172V and G172V+D115N EXO5 exhibit good convergence, with a high Jaccard similarity between replicas, whereas WT EXO5 and L151P EXO5 both display a higher diversity, especially in the case of L151P EXO5.

**Fig. 7.**
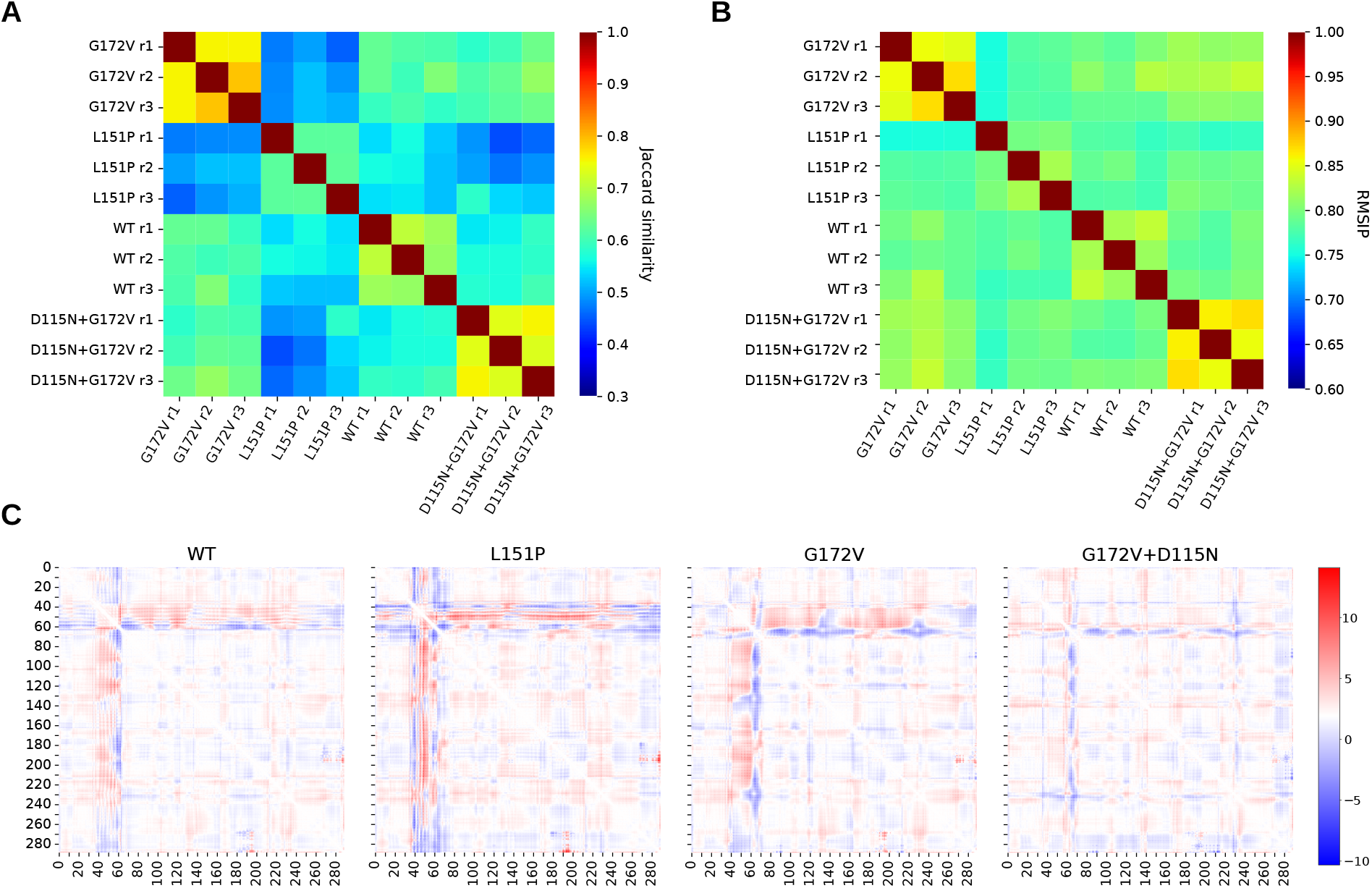
Heatmap of the Jaccard similarity between all unweighted protein structure networks derived from the Pyinteraph analysis, with a 20% cutoff of the occurrence of interactions (A). Root Mean Square Inner Product of the Essential Subspace constituted by the first 30 Principal Components of each of the 3 MD simulation replicas per each EXO5 structure (B). Average change in distance between pairs of aminoacids in the equilibrated trajectories, compared to the starting structures. In red are protein residues that on average across all replica tend to get farther away from each other, in blue residues that on average get closer to each other. Note that residue indices are shifted by -68 compared to the original EXO5, and here they represent the residue index in the simulated structures (C).

Furthermore, the root-mean-square inner product (RMSIP) of the essential subspaces (Figure 7B) shows that G172V and G172V+D115N EXO5, followed by WT EXO5, have the highest similarity in their essential dynamics, highlighting a strong consistency in conformational sampling across different replicas of these variants. In contrast, the RMSIP for L151P EXO5, when compared to other EXO5 variants, is similar to the RMSIP observed within L151P replicas, suggesting that the dynamic differences characteristic of L151P simulations are not readily apparent. The RMSIP values between WT and L151P EXO5 simulations are compatible with the ones between L151P EXO5 and the stable forms of EXO5, suggesting that the essential dynamics captured by Principal Components does not capture any meaningful similarities between the two forms of EXO5 which exhibit the strongest instabilities in the *α*4 region. Figure 7C displays the average pairwise displacement of *α*-carbons from the initial distance map. There is a consistency in the movement of the *α*4 helix away from the core of the protein in all EXO5 forms, as well as an approach of the linker between *α*4 and *α*5 towards *α*5. L151P however shows again a unique behaviour, with residues 106-113 that, on average, tend to lay closer to the rest of the protein. *α*4 itself, on the contrary, moves away from the protein core and thus from DNA in a more pronounced manner with respect to the other EXO5 forms, allowed by its greater degree of disorder.

Overall, both when focusing on local interactions, overall conformational changes, and essential dynamics, all metrics considered show distinct patterns for each EXO5 haplotype, with L151P EXO5 exhibiting the most divergent behaviour.

### Impact of the L151P EXO5 variant on cancer patients

Given the significant structural impact of *haplotype 4* on the EXO5 protein and the previous association of SNP rs35672330 with prostate cancer (PCa) risk characterized in (11), we investigated the distribution of *haplotype 4* in both healthy individuals and cancer patients, utilizing data from the 1000 Genomes Project and TCGA.

Utilizing TCGA genotype and ancestry data obtained from (48) via (50)(62), we reconstructed the EXO5 haplotypes for 8,535 cancer patients spanning 24 cancer types and representing the five major population groups.

No cancer- or population-specific enrichment of *haplotype 4* was observed. Notably, focusing on European individuals, representing the majority of individuals in TCGA data, the frequency of *haplotype 4* was 5.93% in 1000 Genomes Project individuals and 6.18% in TCGA PCa dataset, suggesting that, contrary to (11), SNP rs35672330 may not play a specific role in prostate cancer predisposition.

We hence hypothesized that the L151P EXO5 variant might influence prostate cancer progression and/or aggressiveness. To explore this, we obtained TGCA Overall Survival (OS) and Progression-Free Interval (PFI) data from (52) and stratified TCGA PCa patients based on the presence or absence of *haplotype 4*. We then conducted a Cox proportional hazards regression analysis, adjusting for ancestry. Notably, patients carrying *haplotype 4* on at least one allele exhibited significantly (p-value *<* 0.05) shorter progression-free survival (Figure 8A).

**Fig. 8.**
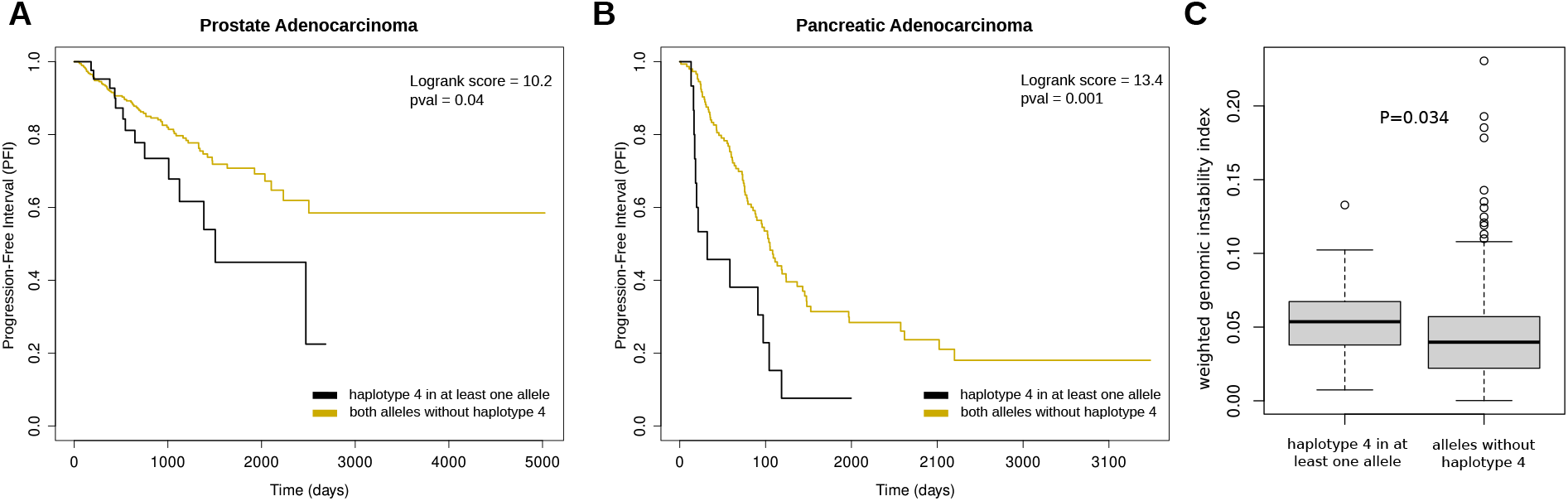
A) Progression-free Interval of patients with prostate cancer stratified by presence of *haplotype 4* in at least one allele; B) Progression-free Interval of patients with pancreatic cancer stratified by presence of *haplotype 4* in at least one allele. C) Weighted genomic instability index values in patients with prostate and pancreatic cancer stratified by presence of *haplotype 4* in at least one allele. Statistic was performed using two-tailed Wilcoxon test statistics.

Intrigued by these findings, we expanded our analysis to include 17 additional cancer types from the TCGA dataset, selecting for the ones comprising more than 100 patients. After correcting for multiple hypothesis, we identified Pancreatic Adenocarcinoma (q-value *<* 0.05) as another cancer type where patients carrying *haplotype 4* on at least one allele show significantly shorter progression-free survival (Figure 8B).

Given the hypothesis that worse progression in these patients could be driven by increased genomic instability, we retrieved genomic instability estimates from (54) and analyzed their association with the presence of *haplotype 4*. As illustrated in Figure 8C, patients carrying *haplotype 4* in at least one allele exhibited a statistically significant increase in genomic instability.

Notably, repeating these analyses while focusing on *haplotypes 1, 2*, or *3* revealed no associations with survival outcomes or genomic instability.

Overall, these findings support the hypothesis that the L151P EXO5 protein variant may contribute to earlier cancer progression across different cancer types by modulating genomic instability.

## Discussion

The study of common human genetic variants offers valuable insights into the biological mechanisms underlying complex traits and diseases. Single nucleotide polymorphisms (SNPs), the most prevalent type of DNA polymorphism, have been shown to influence cancer susceptibility and play a critical role in shaping treatment responses and clinical outcomes, particularly when located in genes encoding proteins involved in DNA damage response (DDR) and repair (4)(5).

In this study, we focused on EXO5, a single-stranded DNA exonuclease involved in DNA repair and previously associated with cancer susceptibility (11)(63). Through an integrated approach that combines large-scale genomic data analysis, protein language models, and molecular dynamics simulations, we investigated the impact of common human EXO5 gene haplotypes on the structural and dynamic properties of the EXO5 protein. Additionally, we explored their potential role in cancer progression using data from comprehensive genomic datasets.

Our results demonstrate a strong correlation between the fitness predicted by protein language models and the maintenance of the structural integrity of the *α*4 helix and its adjacent linkers in EXO5. The importance of this protein region was demonstrated in (12), where it was found that mutations of conserved residues in the *α*4 helix can nearly abolish EXO5’s activity and reduce its binding affinity to ssDNA. The ESM1v model correctly identified the proline substitution Leu151Pro in *α*5 as causing structural instability, which aligns with experimental results obtained in (11) where the substitution was observed to cause a loss of EXO5 nuclease activity and of nuclear localization. Interestingly, it also identified the G172V variant of the EXO5 protein as having a higher fitness than the WT EXO5, an insight that was not immediately explainable from structural considerations alone, but was suggested by our MD simulations to be also related to the stability of helix *α*4.

While our molecular dynamics simulations revealed consistent differences in the *α*4 behavior across systems and replicas, we remark that modeling partially disordered protein regions presents inherent challenges. The Amber ff14sb force field, combined with the TIP3P water model, may introduce some bias in helix propensity predictions (64). However, the robust patterns we observed across different systems suggest that our key findings regarding the structural and dynamic properties of EXO5 variants are reliable. These results align well with both our protein language model predictions and previous experimental observations, providing a multi-faceted view of EXO5 dynamics.

In all systems, the *α*4 region maintains high mobility, but its conformational ensemble and its contacts to the rest of the protein appear to be unique to each EXO5 protein variant. The effects of aminoacid substitutions are often non-local and allosteric in nature, and while in the case of L151P EXO5 we discovered a possible pathway by which the local disruption in *α*5 propagates, in the case of G172V it is unclear from our simulations how the SNP related to a distant loop affects the behavior of *α*4. This highlights the complexity of protein dynamics-function relationship and the importance of considering long-range allosteric effects, even if a substitution does not cause detectable local structural changes.

To understand the biological implications of these structural findings, we focused particularly on the L151P substitution. While it was previously experimentally observed in literature (11), our study provides additional mechanistic insights into its effects on EXO5 dynamics and the potential impact it has on EXO5 nuclear localization. The formation of new stable contacts involving the NLS region in the L151P EXO5 protein variant suggests a possible mechanism for a sharp decrease in nuclear transport binding affinity, which could result in a mostly non-functional EXO5, strongly supporting the findings from (11).

Although it remains unclear whether this fully inactivates EXO5 *in vivo*, the analysis of a large cohort of cancer patients from the TCGA project shows that cancer patients carrying rs35672330 SNP exhibit increased genomic instability and worse prognosis across multiple cancer types. These results, together with the findings by (12), which link EXO5 depletion to increased sister chromatid exchanges—a typical indicator of DNA repair defects or excessive DNA damage—support the hypothesis that the L151P protein variant, by significantly impairing EXO5 function, may exacerbate genomic instability, accelerating disease progression.

In summary, while providing important mechanistic insights on the germline determinants of EXO5 protein structure and dynamics and their possible role in cancer progression, our study also demonstrates the potential of combining large scale genomics data with deep learning variant effect prediction tools and molecular dynamics simulations to explore the impact of common SNPs on protein structure, dynamics and function.

This study primarily focused on the effects of SNPs, leaving *haplotype 5* aside. The latter, being characterized by the presence of an INDEL, introduces a premature stop codon. Hence, as a next step, we plan to integrate the current approach with experimental work to further explore the potential functional consequences of EXO5 germline determinants.

From a technical perspective, sequence-only protein language models like ESM-1v offer distinct advantages for our analysis. While structure-aware methods such as inverse folding (65) and structure-augmented PLMs (66; 67) may achieve higher accuracy on average for structured proteins (68), they might show a systematic accuracy bias toward structured regions, as well as requiring input structures that often require a partial or complete computational determination, possibly introducing additional biases. Fitness prediction using PLMs likelihoods and sequences as the only inputs offers advantages in terms of speed and flexibility compared to many other methods, and while in this work we computed the Pseudo Log-Likelihood of sequences as the sum of log-likelihoods of aminoacids, normalization of the PLL on the sequence length constitutes a natural extension of the scoring method. This allows for the comparison between sequences of different length, as in the case of different isoforms or in the presence of indels, thus enabling the study of complete genetic landscapes. Future work could nonetheless explore the use of other models with different training objectives, different scoring methods, or the use of supervised fine-tuning on experimental variant data. These approaches might allow to distinguish the effects of genetic variants on structural stability and other fitness components.

To conclude, our methodology provides a novel and valuable computational framework to investigate the local interaction of multiple germline variants, which complemented with orthogonal approaches (69)(70), could enhance our ability to explore the interaction landscape of common genetic variants and somatic phenotypes and signatures (71)(72)(73)(74)(75) that underpin cancer risk and progression.

## Data availability

The Molecular Dynamics simulation dataset is accessible on Zenodo at DOI: 10.5281/zenodo.14008334.

## Author contributions

A.R. and G.L. designed and supervised the study. F.M and

D.D. performed the analyses and simulations with inputs from A.B., G.L and A.R.. A.R. and F.M. wrote the manuscript with inputs from D.D, A.B and G.L. All the authors approved the manuscript.

## Funding

The research leading to these results has received funding from the Pezcoller Foundation (PhD fellowship to F.M.) and from Fondazione AIRC under MFAG 2017 - ID. 20621 project - P.I. Alessandro Romanel. A.B. and G.L. acknowledge support from “ICSC – Centro Nazionale di Ricerca in HPC, Big Data and Quantum Computing”, project funded under the National Recovery and Resilience Plan (NRRP), Mission 4 Component 2 Investment 1.4 - Call for tender No. 1031 of 17/06/2022 of Italian Ministry for University and Research funded by the European Union-NextGenerationEU (proj. nr. CN-00000013).

## Acknowledgments

We acknowledge the CINECA award under the ISCRA initiative, for the availability of high-performance computing resources and support. We are grateful to Alberto Inga for critical reading of the manuscript.

## Competing interests

No competing interest is declared.

## Notes

https://zenodo.org/records/14008334

### Competing Interest Statement

The authors have declared no competing interest.

