## Supplementary Information and Figures for "A common haplotype in the EXO5 gene can impact its protein structure and dynamics and modulate genome stability and cancer progression"

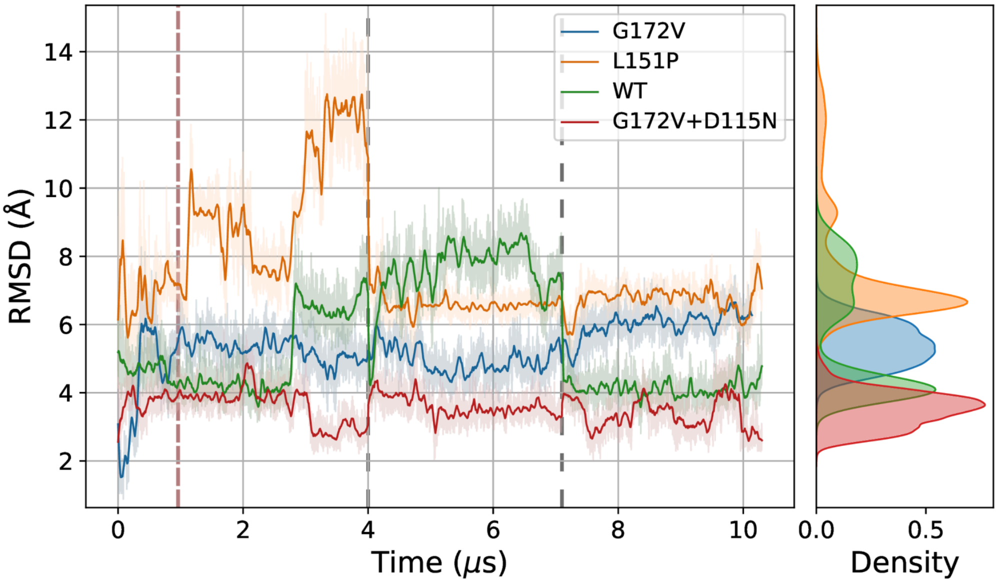


***Figure S1****: RMSD of residues 106-138 along the MD simulations. Frames aligned to the whole initial EXO5 structure. Lighter shades are the raw data, calculated every 0.5 ns, while the darker lines are a moving average with a 40 ns uniform window. The dashed red line indicates the equilibration cutoff, while the dashed gray lines separate different replicas.*

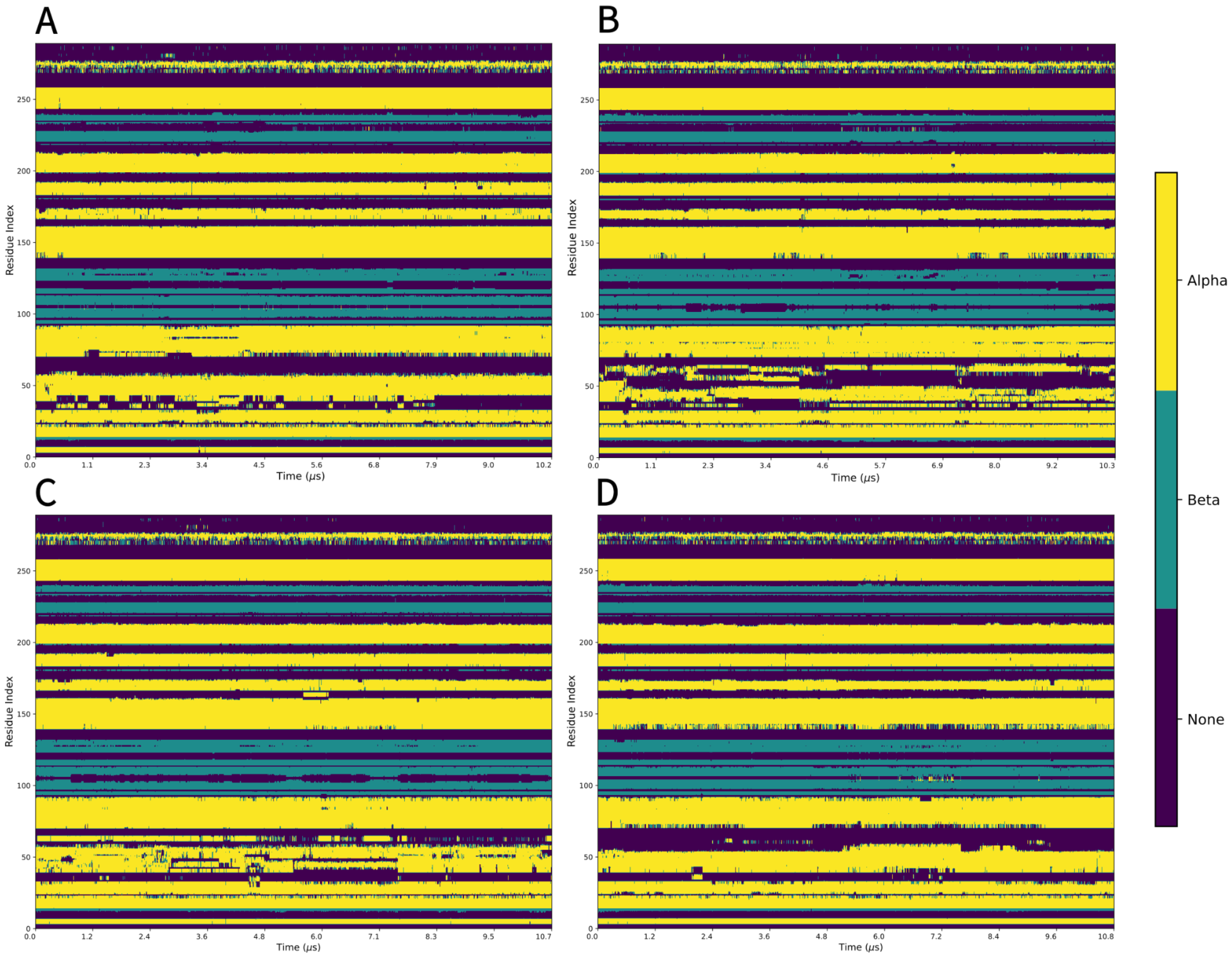


***Figure S2***: *Secondary structure of the EXO5 proteins over the MD simulation replicas, as determined by MDAnalysis using PyDSSP: G172V EXO5 (A), L151P EXO5 (B), WT EXO5 (C), G172V+D115N EXO5 (D). Note that residue indices are shifted by -68 compared to the reference sequence used in the main text.*

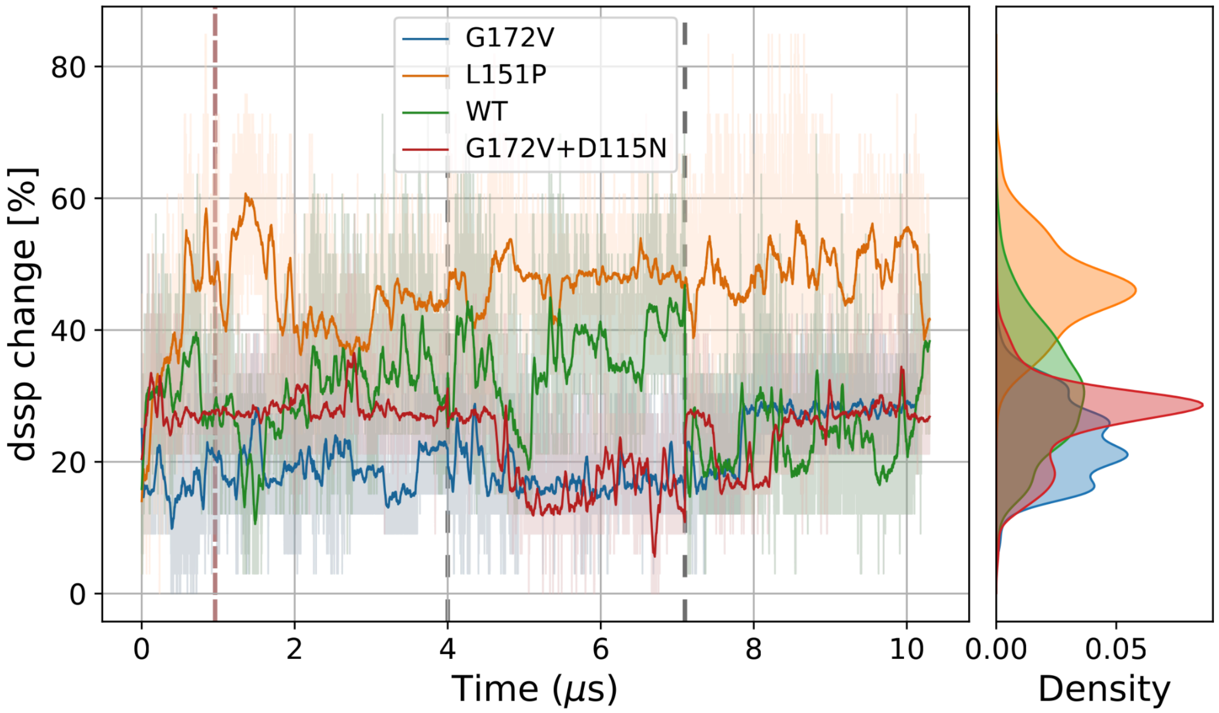


***Figure S3***: *Change in the PyDSSP-assigned secondary structure of the $\alpha4$ region. Lighter shades are the raw data, calculated every \SI{0.5}{\nano\second}, while the darker lines are a moving average with a \SI{40}{\nano\second} uniform window. The dashed red line indicates the chosen equilibration cutoff, while the dashed grey lines separate different replicas.*

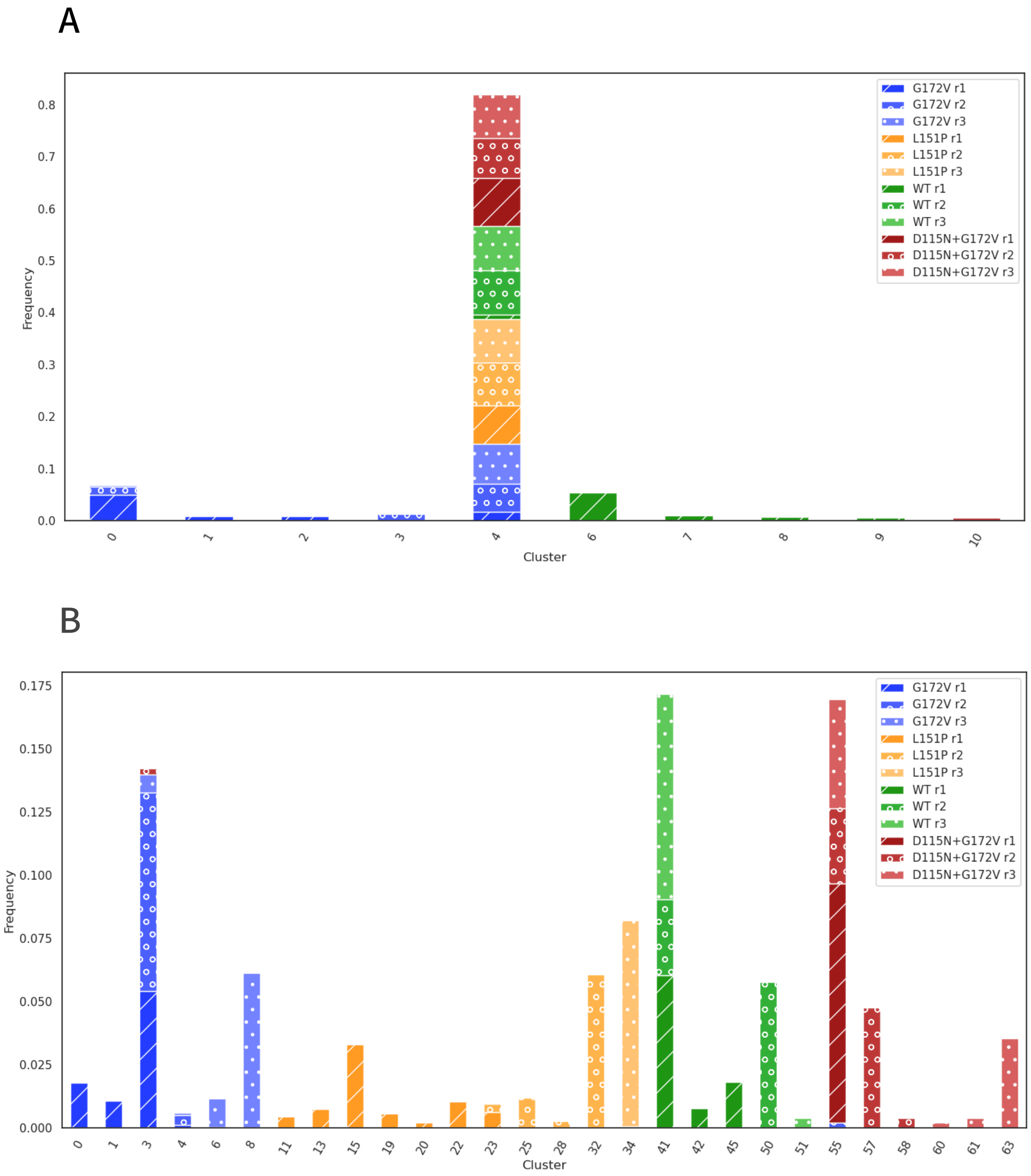


***Figure S4**: Backbone dihedral clustering results of the EXO5 structures excluding the alpha4 region (A) and in the alpha4 region (B). Obtained using Advanced Density Peaks from the DADApy package, with Z=3, halo=False (A) and Z=5 (stricter condition to classify two clusters as separate), halo=False (B). Clusters containing less than 0.5% of frames are not shown. Clusters 3, 8, 15, 32, 34, 41, 50, 55, 57 and 63 are the ones shown in Figure 4B of the main text.*

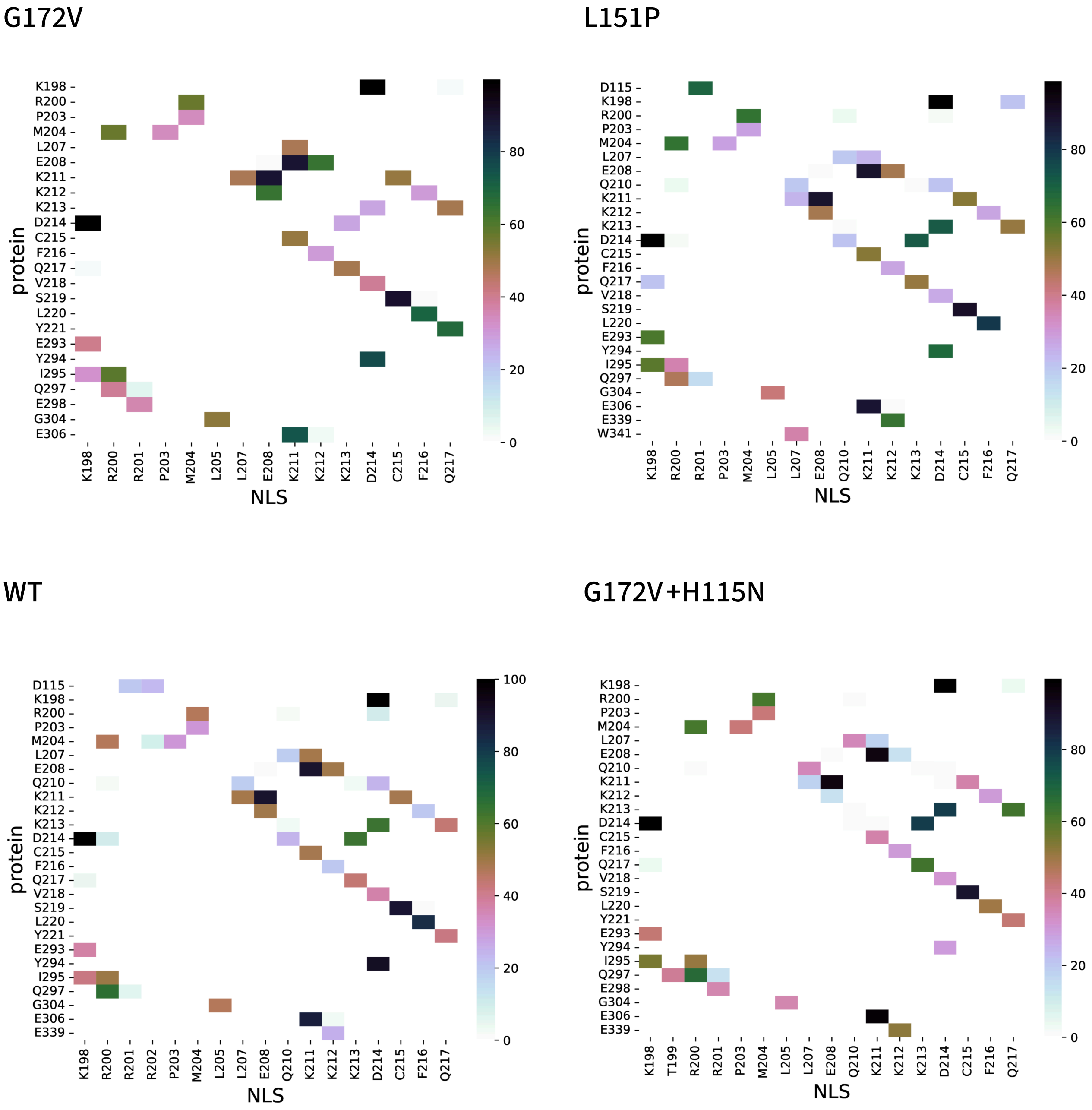


***Figure S5**: Contact maps with the occurrence of interactions between residues of the Nuclear Localization signal and the rest of EXO5, averaged across all replicas. Contacts with less then 20% of occurrence are not shown.*

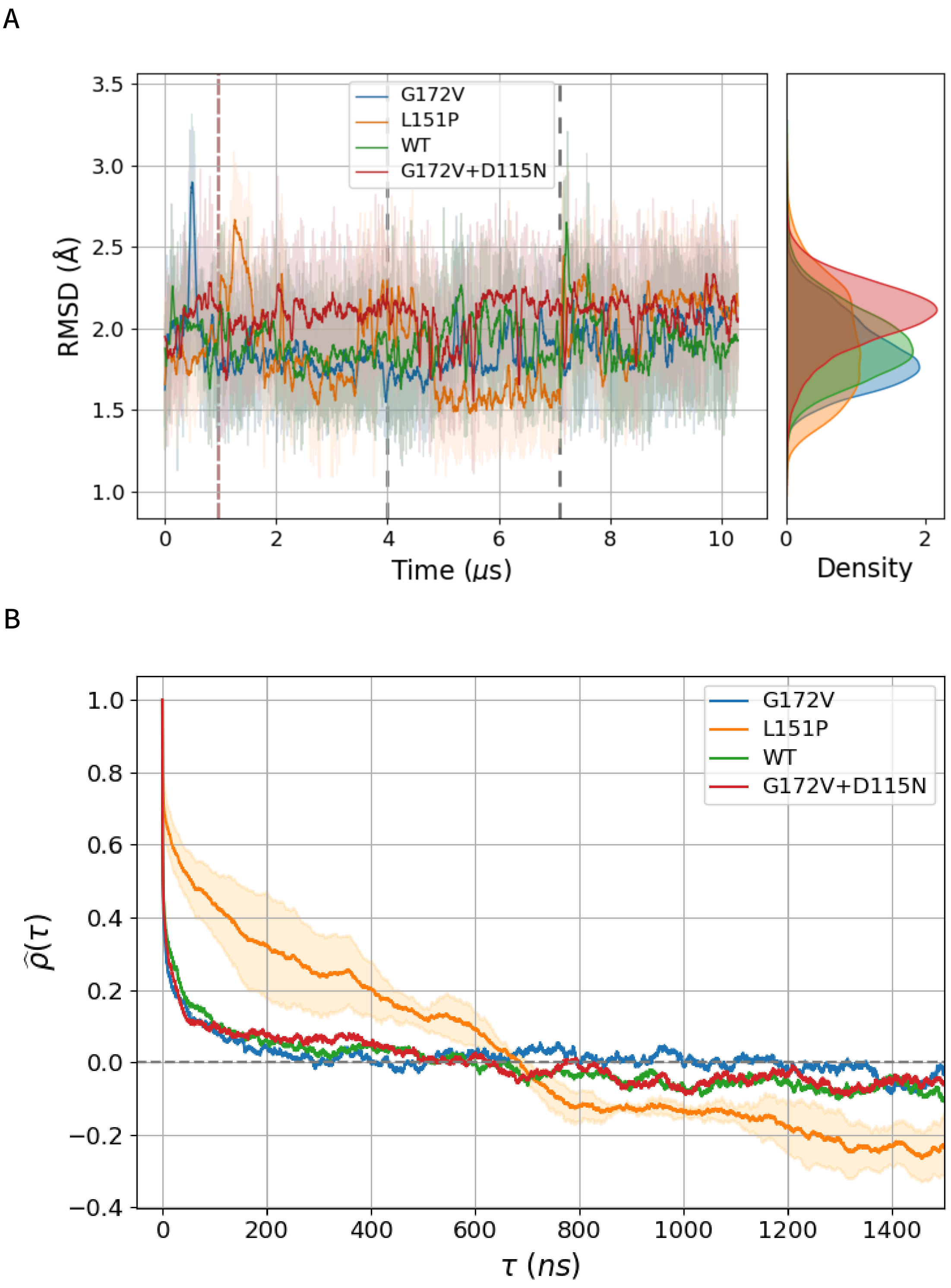


***Figure S6***: *RMSD of the Nuclear Localization Signal (residues 197-215) across all replicas (A). Autocorrelation function of the RMSD of the NLS, obtained by averaging the autocorrelation functions computed separately for each replica (B). For L151P EXO5, the standard error of the mean is shown as the light orange shade. Errors for the other structures are not shown for clarity.*
